# Potent and Broad HIV-1 Neutralization by a Bispecific CD4-CD4i Fusion Protein based on Single-Domain CD4-D1 and X5 CD4i antibody

**DOI:** 10.64898/2026.04.22.719690

**Authors:** Xiaojie Chu, Cory Shetler, Zeyu Sun, Chris Perrone, Zhiwei Feng, Kerri J. Penrose, R. Brad Jones, John W. Mellors, Dimiter S Dimitrov, Wei Li

## Abstract

Human immunodeficiency virus (HIV) infection remains a global health threat. Although antiretroviral therapy (ART) has significantly transformed HIV into a manageable chronic disease, emergence of drug resistance to current ART is a continuing concern. Broadly neutralizing antibodies, as well as reagents containing both a soluble CD4 mimetic and an HIV co-receptor inhibitor—such as CD4-CD4i antibodies—are promising strategies for the prevention and treatment of HIV infection. We previously developed a CD4 D1mimetic (mD1.22) with enhanced neutralization potency, surpassing that of the clinically validated sCD4-D1D2 mimetic. We also previously identified a novel CD4i antibody (X5) that targets the conserved coreceptor binding site on the gp120 core and recognizes an epitope partially overlapping with 17b monoclonal antibody binding site. X5 binding to gp120 was augmented by CD4 and modestly enhanced by CCR5. To leverage these favorable interactions, we designed and optimized sCD4-X5 bispecific antibodies by computational structure-aided modification of antibody size and fusion linkers between the two binding moieties. The bispecific D1X5, with a (G_4_S)_7_ long linker between D1 and X5 (IgG1-LL D1X5), exhibited broad neutralization against 11 diverse HIV subtypes across B, C, G clades and AC, BC recombinants. The TZM-bl cell neutralization assay showed IgG1-LL D1X5 neutralization geometric mean IC50 and IC80 are 0.6 μg/mL and 3.4 μg/mL respectively, which are within the range of potent bnAbs. This work has identified a novel single domain soluble CD4 based CD4-CD4i bispecific antibody with broad HIV-1 neutralization.

## 1. Introduction

Human immunodeficiency virus type 1(HIV-1) is responsible for most of HIV infections, and it remains a significant global health concern. Since the discovery of HIV-1 in 1981, approximately 88.4 million people have acquired the infection. HIV-1 compromises the immune system by targeting and depleting CD4+ T cells, leading to acquired immune deficiency syndrome (AIDS). The advent of antiretroviral therapy (ART) has significantly transformed the disease into a manageable chronic condition. However, resistance to current antiretrovirals remains a continuing threat to ART effectiveness, necessitating new strategies to block HIV-1 replication. Despite these, there is still no cure for HIV infection and a novel therapeutic strategy for a complete cure is urgently needed.

Antibody-based therapeutics that leverage broadly neutralizing antibodies (bNAbs) are a promising approach for the prevention and treatment of HIV-1 infection. It has the potential to provide long-lasting viral suppression, broad spectrum neutralization, and even a functional cure. So far, only one antibody, Ibalizumab-uiyk (Trogarzo), has been approved by the FDA for treating multi-drug resistant HIV-1[1, 2]. It is a CD4-directed post-attachment inhibitor that prevents interaction with the CCR5 and CXCR4 co-receptors, thereby blocking viral entry into CD4+ T cells[3, 4]. Other bnAbs that target vulnerabilities on HIV-1 envelope glycoproteins gp120 and gp41 that facilitate viral attachment and membrane fusion via interaction with CD4 and coreceptors have been reported to suppress HIV-1 replication and control viremia [5-7].

Soluble CD4 (sCD4) and CD4 mimics are attractive HIV-1 entry inhibitors that block HIV-1 infection through multiple mechanisms: i) competitively blocking Env engagement with host cell surface CD4[8, 9]; ii) pre-triggering and inactivation of Env[10]; and iii) exposure of non-neutralizing epitopes and synergizing with non-neutralizing Abs (nnAbs) for mobilization of ADCC[11]. However, sCD4 failed to reduce viremia in clinical trials despite efforts to prolong its half-life by fusion with human IgG Fc fragment (CD4-Ig) and to enhance potency via multivalency constructs such as Pro542[12]. One of the major hurdles for sCD4 is its potential to promote HIV infection of CD4-CCR5+ host cells by inducing a transiently activated state in Env, which adopts an open functionally-competent conformation capable of engaging coreceptors [13-15]. The longevity of this transient fusion competent state depends on intrinsic properties of Env associated with different HIV-1 variants [16].

To prevent sCD4-mediated enhancement of HIV infection, bifunctional molecules that combine sCD4 with additional binding moieties targeting fusion-related domains exposed following sCD4 engagement have emerged as promising strategies. These strategies include targeting the coreceptor binding site; the cluster-A epitope, which corresponds to the gp120 face occluded by gp41 in the trimeric Env; and the exposed groove between N-heptad repeat (NHR) coiled-coil domain of trimeric gp41. CD4-induced (CD4i) antibodies [13] target the conserved HIV-1 coreceptor binding site on gp120, which is exposed following CD4-induced conformational changes. On the monomeric gp120 core, CD4i epitopes are positioned adjacent to the V3 loop base and span the four-stranded bridging antiparallel β-sheet formed by the gp120 β20 and β21 strands, along with the β2 and β3 strands from the V1V2 base. Examples of CD4i antibodies include 17b, 412d, E51, 21c, and 48d [17-22]. Engineering sCD4 and CD4i antibodies into a bifunctional molecule as an HIV entry inhibitor is an attractive strategy due to the spatial proximity between the CD4 binding site and CD4i epitope, the conserved nature of both epitopes, and the potential for synergistic effects of their engagement. Several studies have demonstrated that CD4-CD4i antibodies exhibit enhanced neutralization potency and breadth[23, 24], such as sCD4-17b[24, 25], and the eCD4-Ig, which combines sCD4-D1D2 with the CCR5 N terminal-mimicking sulfated tyrosine peptide[26]. However, translating eCD4-Ig into a protein therapeutic may present manufacturing challenges, particularly due to the requirement for co-transfection or co-expression of sulfotransferases[27]. In non-human primate studies, eCD4-Ig has been administered via AAV-mediated transgene delivery rather than as a purified protein. Currently, most sCD4-CD4i bispecifics including eCD4-Ig are based on the two domain sCD4-D1D2 scaffold. We have successfully developed an engineered cavity-altered monomeric sCD4-D1 domain, mD1.22, which demonstrated higher affinity to gp140 and neutralization potency than sCD4-D1D2[28, 29]. Additionally, mD1.22 is less prone to aggregation than D1D2 and exhibits markedly reduced polyspecificity, likely due to its smaller hydrodynamic radius[30]. Therefore, we aim to reinvigorate the design of CD4-CD4i based on single domain soluble CD4 (D1, mD1.22).

We previously isolated the CD4i antibody X5 from a phage display library, which exhibited high binding affinity against a variety of Envs[18, 31-33]. X5 targets the proximity of the CD4 and coreceptor binding sites on gp120, centered around the I423/K432 residue cluster, with an epitope that partially overlaps with 17b yet remains distinct from the CCR5 footprint[33]. However, X5 alone showed limited neutralization potency and breadth, due to the necessity of CD4 engagement and the spatial restriction nature of the CD4i epitope at the virus-cell interface[32]. It has been shown that the smaller size X5 (scFv and Fab) exhibit higher neutralization potency than the full-length IgG X5[32]. In this study, we designed and constructed a bispecific antibody based on mD1.22 and X5. We hypothesis that tethering mD1.22 to X5 induces gp120 conformational rearrangement *in cis*, thereby bypassing the requirement for cell surface CD4 to expose the X5 epitope and alleviating the spatial constraints that limit X5 neutralization potency. Based in this rationale, we designed and evaluated CD4-CD4i constructs for their neutralization potency across diverse HIV variants and applied structure-guided linker optimization to further enhance their potency.

## 2. Materials and Methods

### 2.1 Expression of bispecific Ab and recombinant gp140 protein

The X5 and mD1.22 (D1) genes were synthesized by IDT and cloned into a pIW-expression vector previously created in our lab[34]. For transient expression of X5-IgG, D1-Fc, D1X5-IgG or scFv, and gp140 (oligomeric gp120-gp41) antigen, the plasmid was transfected into EXPI293 cells by PEI, and proteins were then purified using Nickel resin or Protein A resin, respectively, as previously described[34].

### 2.2 Enzyme-Linked Immunosorbent Assays (ELISAs)

The EC_50_ of bispecific antibodies against recombinant gp140 was analyzed by ELISA as previously described. Recombinant gp140 proteins were coated onto a high binding 96-well plate at 5 μg/ml and incubated at 4°C overnight. The plate was then blocked with 5% MPBS (milk in PBS) for 1 hour at 37°C. After three washes with 0.05% PBST (Tween 20 in PBS), 3-fold serial diluted antibodies were incubated on the plate for another hour at 37°C. Antibody binding was detected using HRP-conjugated goat anti-human IgG1 Fc (Sigma-Aldrich) at a 1:1000 dilution, followed by a 1 hour incubated at 37°C. The plates were washed three times with 0.05% PBST and binding activity was measured by 3,3′,5,5′-tetramethylbenzidine substrate (TMB, Sigma-Aldrich) and was stopped by TMB stop buffer (ScyTek Laboratories). Absorbance was read at 450nm.

### 2.3 Neutralization assay

The antibodies were evaluated for neutralizing activity against a panel of infectious HIV-1 viruses that included: lab-adapted BaL (BEI Resources, NIAID, NIH, ARP-51) and xxLAI [35, 36]; transmitter founder viruses CH0505, CH077, THRO, and REJO; and HIV-1 pseudovirus made using co-transfection of plasmids from the panel of global HIV-1 envelope clones (NIH HIV Reagent Program, NIAID, NIH. HRP-12670, contributed by Dr. David Montefiori) including BJOX2000, X1632, CE1176, 246F3, CH119 with the SG3ΔEnv Non-infectious Molecular Clone (NIH HIV Reagent Program, Division of AIDS, NIAID, NIH, ARP-11051, contributed by Dr. John C. Kappes and Dr. Xiaoyun Wu). HIV-1 BaL was generated by expansion of infectious HIV_BaL_ in activated CD8 depleted PBMCs. All other viruses were made by transfection or co-transfection of plasmids using Lipofectamine 2000 (Thermo fisher) in 293T/17 cells (ATCC) [37].

HIV-1 neutralization was measured in TZM-bl cells (BEI resources, NIADI, NIH, HRP-8129), an indicator cell line derived from HeLa cells that stably express high levels of CD4 and CCR5 and natural CXCR4, and contains luciferase under the control of the HIV-1 promoter and allows quantitative analysis of HIV replicon [38]. Briefly, TZM-bl cells were plated at 10,000 cells per well overnight, then treated with five-fold serial dilutions of antibodies with starting concentrations of 25μg/mL and infected with an HIV-1 variant normalized to an output of 140,000 relative light units (RLU), as determined by endpoint dilution. After a 48-hour incubation at 37°C, cells were lysed, and luminescence was measured in RLU using a commercially available luciferase detection system (BriteLite Plus, Revvity). The 50% and 80% in vitro inhibitory concentrations (IC_50_ and IC_80_) were calculated as the antibody concentration needed to inhibit 50% and 80% of HIV-1 replication in the assay. A batch RPV control virus was included in each experimental setup[37]. Neutralization assays were performed in triplicate for each HIV-1 variant.

### 2.4 Biolayer interferometry (BLItz)

The affinity and avidity of bispecific antibodies were measured by the biolayer interferometry BLItz (ForteBio, Menlo Park, CA), as previously described [34]. Briefly, biotinylated recombinant gp140 protein was coated onto streptavidin biosensors (ForteBio) at 16.7 μg/mL for 2 minutes and equilibrated with Dulbecco’s phosphate-buffered saline (DPBS) (pH = 7.4) to establish baselines. Antibodies at various concentrations were loaded for association monitoring for 2 minutes, followed by a 4-minute dissociation phase in Dulbecco’s Phosphate-Buffered Saline (DPBS). Antigen-coated biosensors incubated with DPBS served as the reference control.

### 2.5 Size Exclusion Chromatography (SEC)

Antibody structure and purity were analyzed using Superdex 200 Increase 10/300 GL chromatography as previously described[34]. A total of 200 μg of filtered D1X5-IgG, D1X5-IgG-LL, and D1X5-scFv-Fc antibodies were analyzed by the ÄKTA explorer machine. Proteins were eluted by DPBS buffer at a flow rate of 0.5 mL/min. Ferritin, Aldolase, Conalbumin, Ovalbumin, Carbonic anhydrase and Ribonuclease were used as the standard proteins for calibration.

### 2.6 Bispecific Antibody Structure Construction and Molecular Dynamic Simulation

The structure of the mD1.22 antibody was generated by homology modeling using SWISS-MODEL[39, 40], with PDB ID: 2NY4[41] automatically selected as the template. Based on the published structure of D1 and X5 in complex with gp120 (PDB ID: 2B4C)[31], missing residues were modeled using the MODELLER[42] “AutoModel” function to construct linker variants of different lengths.

To set up the simulation system, we followed same force fields or parameters as we described before[43, 44]. Specifically, based on the modeled structures, we set a periodic boundary conditions cubic solvation box with the buffer distance of 10 angstrom. Water molecules (TIP3P model) were applied and 0.15M of NaCl was added to system. AMBER22[45] software package was then deployed to execute the simulation.

The MD system was first relaxed through a series of energy minimizations to remove potential steric clashes. This was followed by three distinct phases of NPT simulations, where the number of particles, pressure, and temperature were kept constant. The first phase was a relaxation stage, involving 0.2 nanoseconds of simulation at each temperature increment from 50 Kelvin to 250 Kelvin in steps of 50 Kelvin. The second phase was a 5-nanosecond equilibration stage conducted at 298 Kelvin. Finally, the third phase was a 500-nanosecond sampling stage. The equations of motion were integrated using a time step of 1 femtosecond during the relaxation phase, and 2 femtoseconds during both the equilibration and sampling phases.

From the sampling phase, 100 snapshots were uniformly extracted and subjected to binding free energy decomposition analysis using the Molecular Mechanics/Generalized Born Surface Area (MM/GBSA) method[46]. For each MD snapshot, both the molecular mechanical energy (EMM) and the solvation free energy were calculated using MM/GBSA.

### 2.5 Pharmacokinetic Assessments

BALB/c mice (6-8 weeks old, Jackson Laboratory) were randomly divided into groups (n=3) and intravenous (i.v.) injected with D1X5, X5, D1X5-IgG-LL, or D1X5-scFv-Fc at a dose of 25 mg/kg. Blood samples (10 μL per mouse) were collected at 1h, 4h, 8h, 1d, 2d, 4d, 6d, and 8d post-injection, and serum was diluted 200-fold with PBS. Antibody concentrations in serum were measured by ELISA using binding to recombinant gp140 as the detection readout. Briefly, recombinant gp140 was coated onto a high-binding 96-well plate at 5 μg/ml and incubated at 4°C overnight. The plate was then blocked with 5% BSA at 37°C for 1 hour. After three washes with PBST (0.05% Tween-20), diluted serum was added to each well and incubated at 37°C for 1 hour. The plates were washed three times before adding HRP-conjugated anti-human Fc antibody, followed by an additional 1-hour incubation. After a final wash, TMB (Sigma-Aldrich) was added to develop color, and the reaction were stopped by adding TMB stop buffer. Antibody concentration was calculated based on the standard curve made by D1X5, X5, D1X5-IgG-LL, or D1X5-scFv-Fc protein separately.

### 2.6 Statistical analysis

Statistical analysis including IC_50_ were performed by GraphPad Prism. Significance was tested using unpaired t test. Differences were considered statically significant at *p*< 0.05.

## 3. Result and Discussion

### 3.1 D1X5 bispecific antibody exhibited higher neutralization potency against HIV pseudoviruses than either individual antibodies or their combination

We previously identified the X5 antibody, which efficiently binds a variety of gp120 molecules complexed with CD4 and CCR5 receptors [47]. However, its epitope requires induction and exposure by CD4. To further enhance the neutralization efficiency of X5, we designed a bispecific molecule by fusing the monomeric CD4-D1 domain, mD1.22 (D1) [28, 29], to the N terminus of the X5 IgG1 antibody heavy chain using a (G_4_S)_3_ linker (**Figure 1A**). After expression, the IgG1 D1X5 displayed high purity (**Figure S1**). This bispecific antibody IgG1 D1X5 exhibited the same EC_50_ (0.06 nM) for binding to gp140 by ELISA as the single antibody combination D1-Fc + IgG1 X5. Additionally, this bispecific D1X5 antibody demonstrated a higher binding affinity (EC_50_) than either D1-Fc (0.2nM) or IgG1 X5 (1.1nM) alone (**Figure 1B**). The equilibrium dissociation constants (KD) of IgG1 D1X5 was 0.02nM as measured by BLItz, which was more than 200-fold lower (indicating stronger binding) than D1-Fc (4.5nM) and IgG1 X5 (8nM) (**Table 1 and Figure S2**). IgG1 X5, D1-Fc, and IgG1 D1X5 exhibited monomeric folding as measured by size-exclusion chromatography (**Figure 1C**). The higher affinity of IgG1 D1X5 compared to D1-Fc, combined with their monomeric nature indicates simultaneous binding of two targeting moieties (D1 and X5) to Env CD4bs and CoRbs. The HIV neutralization IC_50_ values of IgG1 D1X5, the combination of D1-Fc + IgG1 X5 added as separate entities, IgG1 X5, and D1-Fc alone against Tier 2 HIV pseudoviruses PSV_p246F3 env_ were 3.1 μg/mL, 9.2 μg/mL, 287.6 μg/mL, and 8.9 μg/mL respectively. The neutralization IC_50_ values of IgG1 D1X5 and D1-Fc against HIV PSV_CH119 env_ were 5.6 μg/mL and 15.1 μg/mL respectively, while the D1-Fc + IgG1 X5 combination and IgG1 X5 alone showed IC_50_ values exceeding 100 μg/mL (**Figure 1D**). These results indicate that the bispecific IgG1 D1X5 antibody exhibits more potent neutralization activity against Tier 2 HIV pseudoviruses compared to single antibodies or antibody combinations.

**Table 1.** The equilibrium dissociation constants (K_D_) of HIV bispecific antibody tested by Blitz.

| Antibody | k<sub>on</sub> (M<sup>-1</sup>s<sup>-1</sup>)<sup>1</sup> | k<sub>off</sub> (s<sup>-1</sup>)<sup>1</sup> | K<sub>D</sub> (nM)<sup>1</sup> |
| --- | --- | --- | --- |
| IgG1 X5 | 6.1 × 10<sup>4</sup> ± 6.2 × 10<sup>2</sup> | 4.9 × 10<sup>-4</sup> ± 2.0 × 10<sup>-5</sup> | 8.0 |
| D1-Fc | 4.1 × 10<sup>4</sup> ± 3.7 × 10<sup>2</sup> | 1.9 × 10<sup>-3</sup> ± 1.4 × 10<sup>-5</sup> | 4.5 |
| IgG1 D1X5 | 5.8 × 10<sup>4</sup> ± 5.5 × 10<sup>2</sup> | 9.4 × 10<sup>-7</sup> ± 1.4 × 10<sup>-5</sup> | 0.02 |
| scFv X5 | 4.5 × 10<sup>4</sup> ± 5.9 × 10<sup>2</sup> | 2.4 × 10<sup>-3</sup> ± 4.3 × 10<sup>-5</sup> | 54.1 |
| scFv D1X5 | 3.3 × 10<sup>4</sup> ± 2.8 × 10<sup>2</sup> | 4.1 × 10<sup>-4</sup> ± 1.4 × 10<sup>-5</sup> | 12.5 |
| scFv-Fc D1X5 | 7.4 × 10<sup>4</sup> ± 6.2 × 10<sup>2</sup> | 2.1 × 10<sup>-4</sup> ± 1.4 × 10<sup>-5</sup> | 2.8 |
| IgG1-LL D1X5 | 4.4 × 10<sup>4</sup> ± 3.6 × 10<sup>2</sup> | 6.7 × 10<sup>-5</sup> ± 1.2 × 10<sup>-5</sup> | 1.5 |

<sup>1</sup>Mean kinetic rate constants ( $k_{on}$ , $k_{off}$ ) and equilibrium dissociation constants ( $K_D = k_{off}/k_{on}$ ) were determined from curve fitting analyses of BLItz sensorgrams.

**Figure 1.**
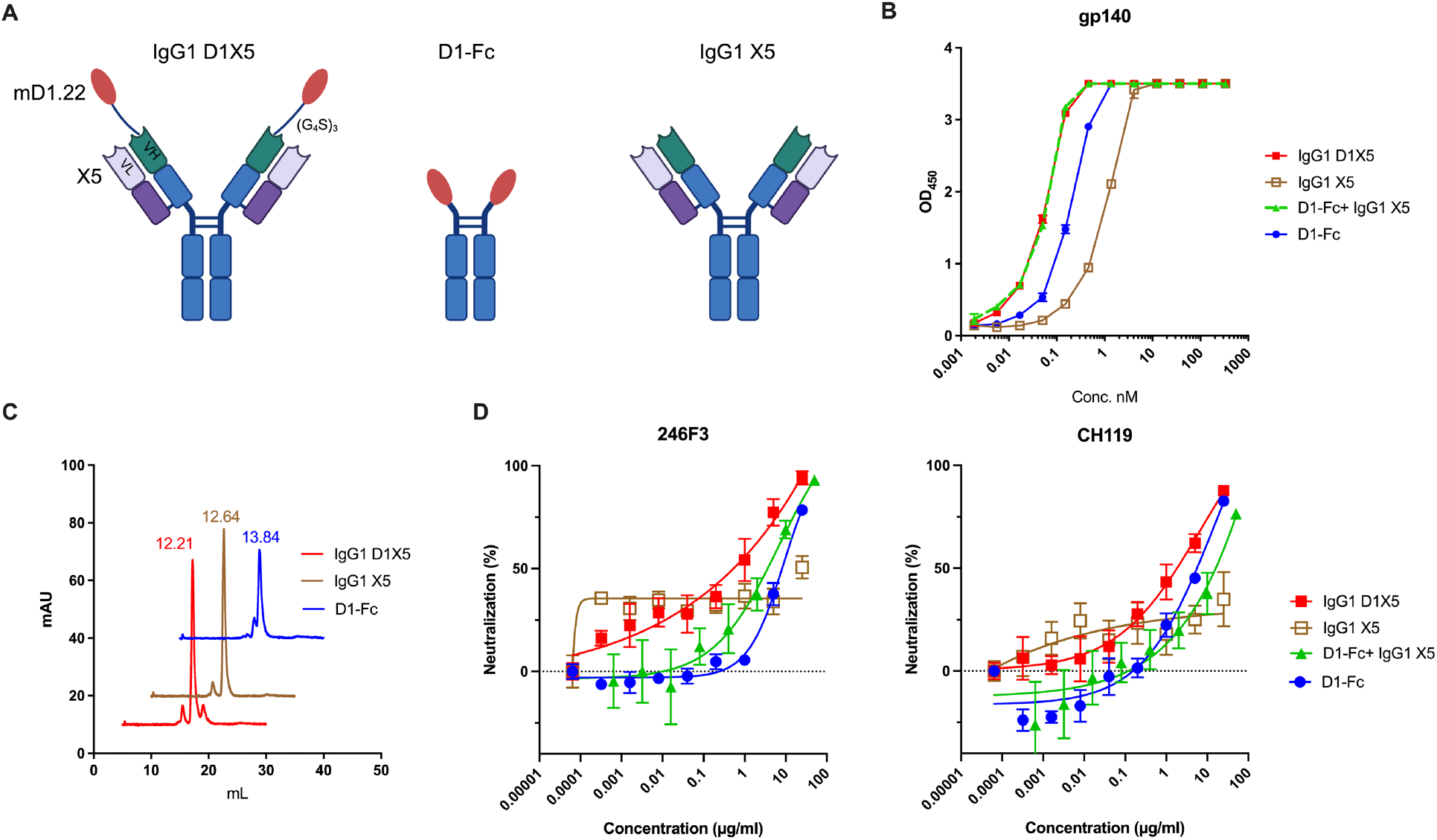
Bispecific IgG1 D1X5 antibody shows enhanced neutralization potency compared with single or combination of antibodies. (A) Schematic design of IgG1 D1X5, D1-Fc, IgG1 X5. The schematic figure was created with BioRender.com (B) The EC_50_ of IgG1 D1X5, D1-Fc, IgG1 X5, and D1-Fc + IgG1 X5 binding to HIV-1 gp140 measured by ELISA. Experiments were performed in duplicate. (C) Aggregation evaluation of IgG1 D1X5, D1-Fc, IgG1 X5 measured by SEC. (D) Neutralization test of IgG1 D1X5, D1-Fc, IgG1 X5, and D1-Fc + IgG1 X5 against HIV-1 PSV variants HIV-1_246F3_ and HIV-1_CH119_. The concentration of D1-Fc in the D1-Fc + IgG1 X5 combination was set at 25 nM; IgG1 X5 was added after 2-fold serial dilution starting from 50 µg/ml. The concentrations of IgG1 D1X5, D1-Fc, and IgG1 X5 as single agents were tested after 2-fold serial dilution, starting at 25 µg/ml. Experiments were performed in triplicate.

### 3.2 Profiling of HIV neutralization potency and breadth for the bispecific IgG1 D1X5 on diverse HIV variants

To investigate the potency and breadth of neutralization activity of D1X5, diverse HIV variants including HIV-1 pseudoviruses (PSVs) from different subtypes, along with Tier 1b or 1a lab-adapted live viruses and Tier 2 full-length transmitted/founder (T/F) viruses infectious molecular clone (IMC) were used for neutralization. Interestingly, all tested panel were insensitive to IgG1 X5 alone (**Figure 2 and Table 2**), which is consistent to the low neutralization capacity for the large-sized full length IgG1 X5. The D1-Fc neutralized most of these strains with varying IC_50_ and IC_80_ ranging from 0.1 to 15.1 μg/mL and 0.3 to 246.6 μg/mL separately, except the T/F IMC clade B CH077 and THRO (IC50 >100 μg/mL). The differential sensitivity of HIV-1 strains to sCD4 reflects either the accessibility of the Env CD4 binding site modulated by surrounding loops and glycan shields or the intrinsic triggerability of Env following CD4 engagement [48-51]. Encouragingly, incorporating X5 into D1-Fc (IgG1 D1X5) moderately improved neutralization potency, reducing the geometric mean IC_50_ from 2.6 μg/mL for D1-Fc to 1.3 μg/mL for IgG1 D1X5, and the IC_80_ from 11.2 μg/mL to 8.9 μg/mL. When normalized to molar concentration by accounting for molecular weight, the neutralization potency of IgG1 D1X5 improved by ~3 folds compared with D1-Fc (IC_50_: 7.2 nM vs. 32.5 nM; IC_80_: 49.5 nM vs. 140.1 nM). Interestingly, for several strains including PSV_CE1176 env_, PSV_246F3_ env, the T/F IMC HIV_CH0505_ and the lab-adapted viruses HIV_Bal_ and HIV_xxLAI_, IgG1 D1X5 exhibited reduced neutralization potency compared with D1-Fc based on IC_80_ values. This reduction is likely due to steric hindrance arising from the larger molecular size of IgG1 D1X5 when approaching the Env CD4 binding site. Impressively, IgG1 D1X5 converted D1-Fc insensitive variants into sensitive ones including T/F IMC clade B HIV_CH077_ and HIV_THRO_), highlighting the synergistic role between X5 and D1.

**Table 2.** Neutralization (IC_50_ and IC_80_, μg/mL) of bispecific antibody against HIV variants.

| HIV Variants | Tier | Subtype | IC50 |  |  |  |  | IC80 |  |  |  |
| --- | --- | --- | --- | --- | --- | --- | --- | --- | --- | --- | --- |
|  |  |  | IgG1<br>X5 | D1-Fc | IgG1<br>D1X5 | IgG1-LL<br>D1X5 | scFv-Fc<br>D1X5 | D1-Fc | IgG1<br>D1X5 | IgG1-LL<br>D1X5 | scFv-Fc<br>D1X5 |
| HIV-1 PSV1 |  |  |  |  |  |  |  |  |  |  |  |
| BJOX2000 | 2 | CRF07_BC | >100 | 0.3 | 0.3 | 0.3 | 0.3 | 2.1 | 1.9 | 2.3 | 2.5 |
| X1632 | 2 | G | >100 | 0.2 | 0.2 | 0.1 | 0.2 | 0.9 | 1 | 0.5 | 1.2 |
| CE1176 | 2 | C | >100 | 2.3 | 2.6 | 0.95 | 0.6 | 18.3 | 42 | 4.2 | 3.3 |
| 246F3 | 2 | AC recomb | >100 | 8.9 | 3.1 | 1.1 | 2.4 | 35.8 | 40.7 | 16.8 | 60.8 |
| CH119 | 2 | CRF07_BC | >100 | 15.1 | 5.6 | 1.6 | 12.2 | 246.6 | 121.8 | 18.3 | 518.6 |
| Lab-adapted strains |  |  |  |  |  |  |  |  |  |  |  |
| Bal | 1b | B | >100 | 0.1 | 0.4 | 0.08 | 0.04 | 0.3 | 0.9 | 0.3 | 0.8 |
| xxLAI | 1a | B | >100 | 0.7 | 0.96 | 0.2 | 0.1 | 1.8 | 2.8 | 0.9 | 0.8 |
| HIV-1 Infectious molecular clone (IMC) Transmitted /founder (T/F) |  |  |  |  |  |  |  |  |  |  |  |
| CH0505 | 2 | C | >100 | 2.2 | 6.8 | 1.1 | 0.9 | 17.3 | 60.3 | 4.1 | 4.3 |
| CH077 | 2 | B | >100 | >100 | 0.7 | 0.7 | 0.4 | >100 | 2.6 | 2.9 | 2.6 |
| THRO | 2 | B | >100 | >100 | 8 | 11.5 | 5.9 | >100 | 120.8 | 71.7 | 33.3 |
| REJO | 2 | B | >100 | 1.4 | 0.4 | 0.5 | 0.4 | 12.6 | 1.5 | 2 | 1.8 |
| Geometric Mean (µg/mL) |  |  | >100 | 2.6 | 1.3 | 0.6 | 0.6 | 11.2 | 8.9 | 3.4 | 5.5 |
| Geometric Mean (nM) |  |  | >667 | 32.5 | 7.2 | 3.2 | 4.1 | 140.1 | 49.5 | 17.9 | 37.9 |

| IC Range |  |  |  |  |  |  |
| --- | --- | --- | --- | --- | --- | --- |
|  | < 0.1 | 0.1 - 1 | 1 - 10 | 10 - 50 | 50 - 100 | > 100 |

<sup>1</sup>PSV: Pseudovirus; IC<sub>50</sub> and IC<sub>80</sub> were in µg/mL. Experiments were performed in triplicate.

**Figure 2.**
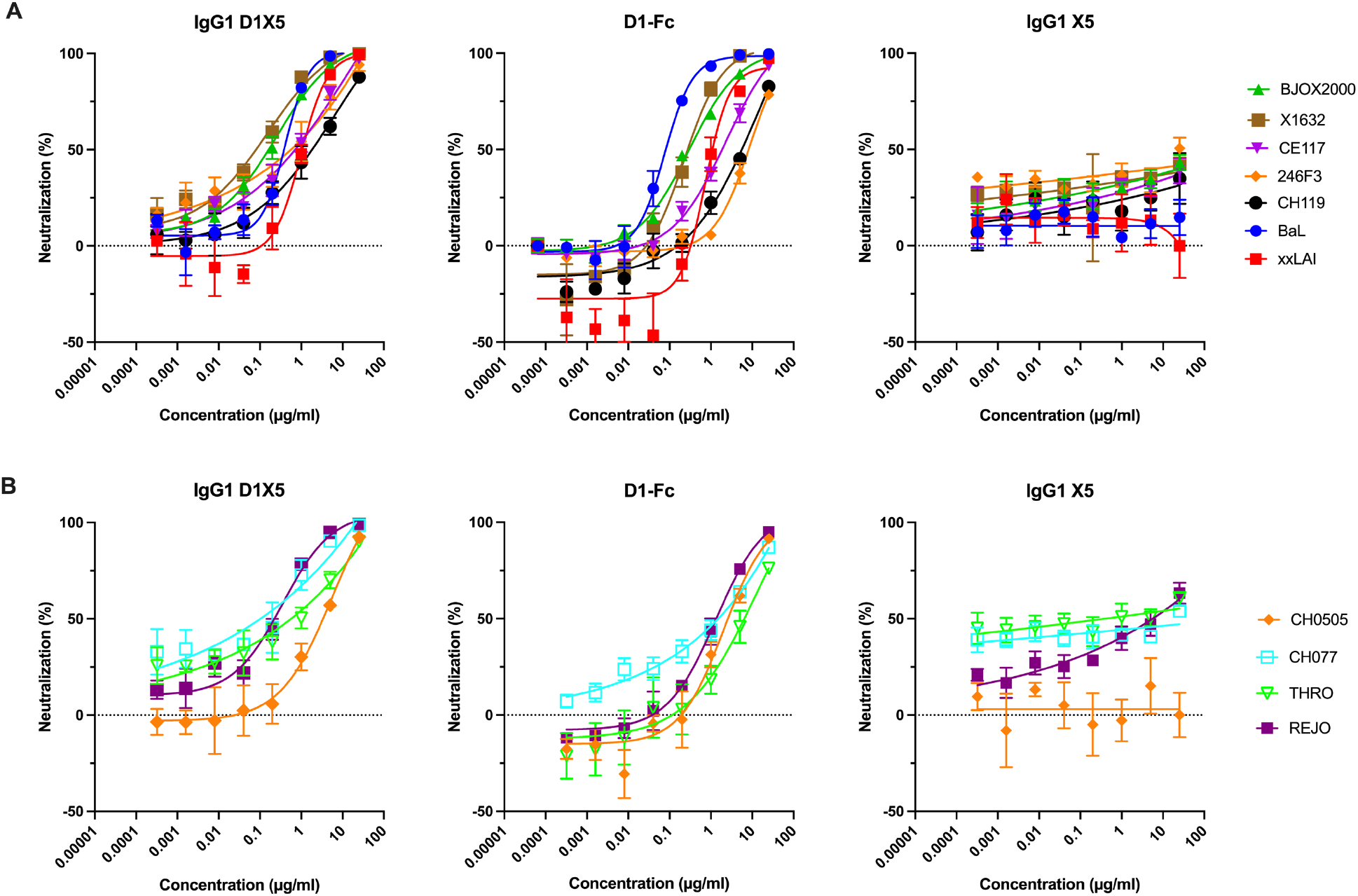
Bispecific IgG1 D1X5 antibody shows broadly neutralization activity. (A) The neutralization by IgG1 D1X5, D1-Fc, IgG1 X5 of HIV-1 PSV variants BJOX2000, X1632, CE117, 246F3, CH119 and lab-adapted variants BaL and xxLAI. (B) The neutralization by IgG1 D1X5, D1-Fc, IgG1 X5 of HIV-1 IMC T/F variants CH0505, CH077, THRO, REJO. Experiments were performed in triplicate.

Overall, IgG1 D1X5 broadly neutralized all tested HIV variants spanning diverse genetic subtypes including clades B, C, and G as well as AC and BC recombinants, with moderate potency, achieving a geometric mean IC_50_ of 1.3 μg/mL and an IC_80_ of 8.9 μg/mL. On this limited neutralization panel, IgG1 D1X5 appeared broadly active, but its potency was lower than that of elite bnAbs, which typically achieve IC_50_ values below 0.5 μg/mL with more than 80% breadth[52].These findings indicate the need to further enhance the neutralization potency of IgG1 D1X5.

### 3.3 Improving neutralization potency through molecular size and linker optimization in the D1X5 bispecifics

X5 is a CD4-induced (CD4i) antibody with spatial restricted epitopes, which is surrounded by V1/V2 and V3 loops. Previous studies have shown that the IgG format of X5 is less potent than the single-chain variable fragment (scFv) format, primarily due to steric constraints limit the ability of X5 to access its gp120 epitope[32, 47]. Therefore, we hypothesized that either reducing antibody size or relieving the steric constrain between D1 and X5 by optimizing the polypeptide linker between them may facilitate simultaneous binding to their recessed epitopes, thereby improving the neutralization potency of D1X5 bispecifics. Based on this hypothesis, we engineered the X5 scFv format and extended the linker length between D1 and X5 from three repeats of (G_4_S) to seven repeats (G_4_S)_7_. To enhance half-life, we engineered both scFv-Fc and IgG formats with a (G_4_S)_7_ long linker between D1 and X5 (**Figure 3A**).

**Figure 3.**
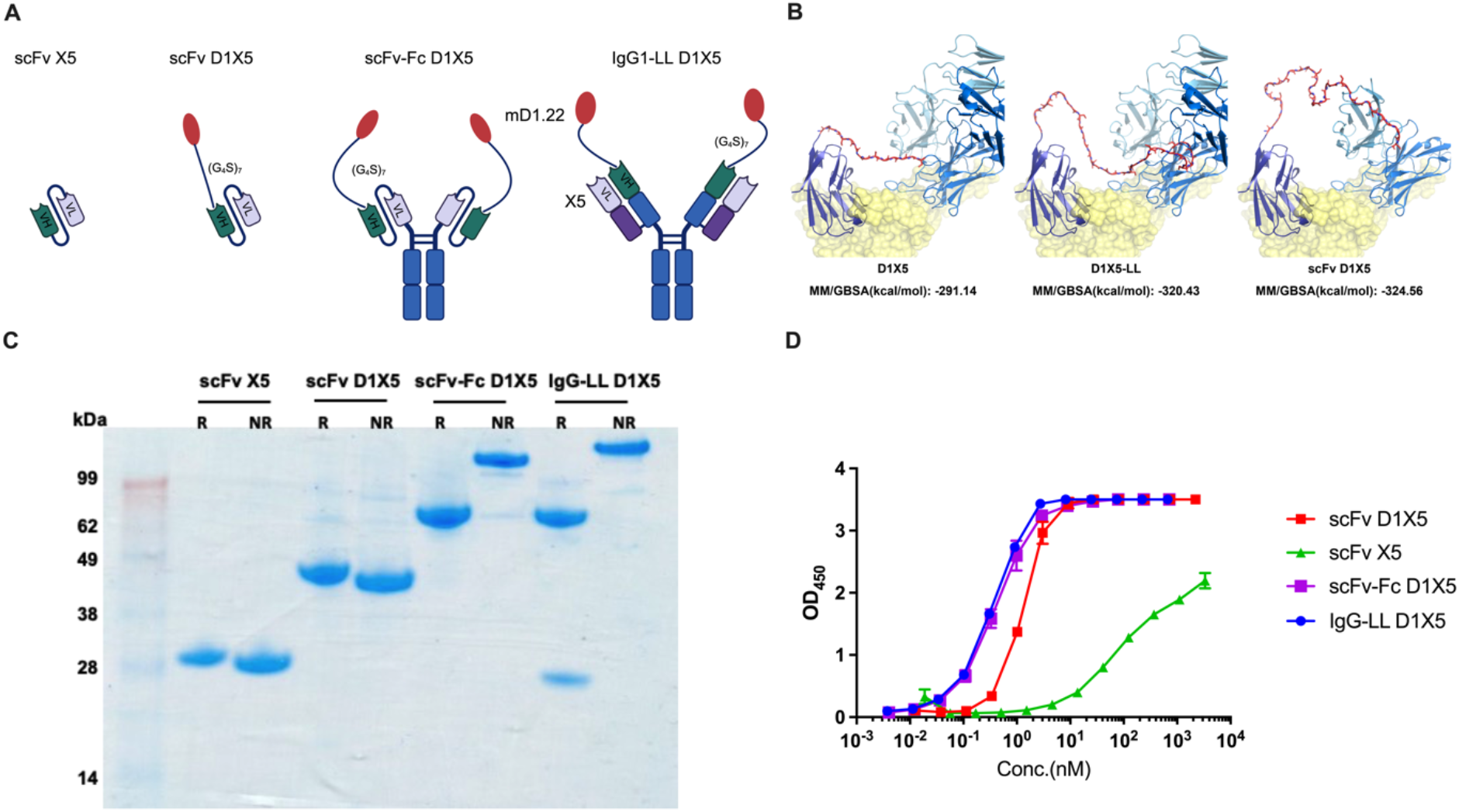
Design and characterization of bispecific D1X5 antibody to enhance HIV neutralization. (A) Schematic design featuring a long linker (G_4_S)_7_ for bispecific scFv D1X5, scFv-Fc D1X5, IgG1-LL D1X5, and the scFv format of X5. The schematic figure was created with BioRender.com (B) In silico binding energy analysis of gp140 for bispecific Fab D1X5 with a short linker (G_4_S)_3_ (left), Fab D1X5 with a long linker (LL) (G_4_S)_7_ (middle), and scFv D1X5 with a long linker (right). (C) The purity validation of scFv X5, scFv D1X5, scFv-Fc D1X5, and IgG1-LL D1X5 by SDS-PAGE stained with Coomassie blue. NR: Non-reducing; R: Reducing. (D) scFv X5, scFv D1X5, scFv-Fc D1X5, and IgG1-LL D1X5 binding to HIV-1 gp140 measured by ELISA. Experiments were performed in duplicate.

The linker length designs were evaluated using the MM/GBSA method. Molecular dynamics (MD) simulations were performed for 500ns, followed by binding free energy calculation. Fab D1X5-LL and scFv D1X5 with (G_4_S)_7_ long linker exhibited lower binding energies to gp120 (−320.43 kcal/mol and −324.56 kcal/mol, respectively) compared to Fab D1X5 with (G_4_S)_3_ short linker (−291.14 kcal/mol). These results suggest that a longer linker might positively contribute to D1X5 binding stability, likely by providing greater conformational freedom to the D1 and X5 moieties to circumvent steric hindrance to reach CD4bs and CD4i epitopes on Env (**Figure 3B**). After expression in mammalian cells, the purity of these D1X5 bispecifics was confirmed to be above 95% (**Figure 3C**). The EC_50_ of scFv D1X5 was highly improved compared to scFv X5 alone (1.3 nM vs. 104.4 nM), suggesting the long (G4S)_7_ linker may allow for simultaneous binding to Env. The EC_50_ values for scFv-Fc D1X5 and IgG-LL D1X5 were 0.4 nM and 0.35 nM, respectively (**Figure 3D**). The equilibrium dissociation constants (KD) of scFv X5, scFv D1X5, scFv-Fc D1X5, and IgG-LL D1X5 were 54.1 nM, 12.5nM, 2.8nM and 1.5nM, respectively (**Table 1 and Figure S2**).

The neutralization potency of scFv-Fc D1X5 were similar to those of scFv D1X5, but both were improved compared to scFv X5 alone (**Figure S3 and Table S1**). The potency of scFv-Fc D1X5 was enhanced compared with D1-Fc (geometric mean IC_50_: 0.6 vs. 2.6 μg/mL; IC_80_: 5.5 vs. 11.2 μg/mL; or in molar units: 4.1 vs. 32.5 nM for IC_50_ and 37.9 vs. 140.1 nM for IC_80_), reaffirming the synergistic effects of X5 and D1. The potency of scFv-Fc D1X5 was moderately improved compared with IgG1 D1X5 (geometric mean IC_50_: 0.6 vs. 1.3 μg/mL; IC_80_: 5.5 vs. 8.9 μg/mL; or in molar units: 4.1 vs.7.2 nM for IC_50_; 37.9 vs. 49.5 nM for IC_80_). The most striking improvement in neutralization was observed with IgG1-LL D1X5, which achieved a geometric mean IC_50_ of 0.6 μg/mL and an IC_80_ of 3.4 μg/mL, representing more than a 4-fold reduction compared with D1-Fc (IC_50_: 2.6 μg/mL; IC_80_: 11.2 μg/ml) and a 2-fold reduction compared with IgG1-D1X5 (IC_50_: 1.3 μg/mL; IC_80_: 8.9 μg/ml). The median IC_50_ and IC_80_ values of IgG1-LL D1X5 were also lower than those of D1-Fc (unpaired t test: p= 0.12 for IC_50_, P=0.13 for IC_80_) and IgG1-D1X5 (p= 0.46 for IC_50_, P=0.13 for IC_80_) (**Figure 4B**). Although these differences did not reach statistical significance, the p-value for IC_80_ were substantially reduced when the linker length was extended in IgG D1X5-LL and scFv-Fc D1X5 compared with IgG D1X5 relative to D1-Fc. (p=0.13 vs. p=0.64), suggesting a trend toward improved potency with longer linkers. These results indicated that the (G_4_S)_7_ long linker facilitates HIV neutralization, consistent with the computational predictions. As scFv-Fc D1X5 incorporates both the (G_4_S)_7_ long linker and the smaller sized X5, its geometric mean IC_80_ nevertheless remained higher than that of IgG1-LL D1X5 (5.5 vs. 3.4 μg/mL), indicating that the enhanced HIV neutralization is driven primarily by the extended (G_4_S)_7_ linker rather than the reduced size of X5 (**Figure 4B**).

**Figure 4.**
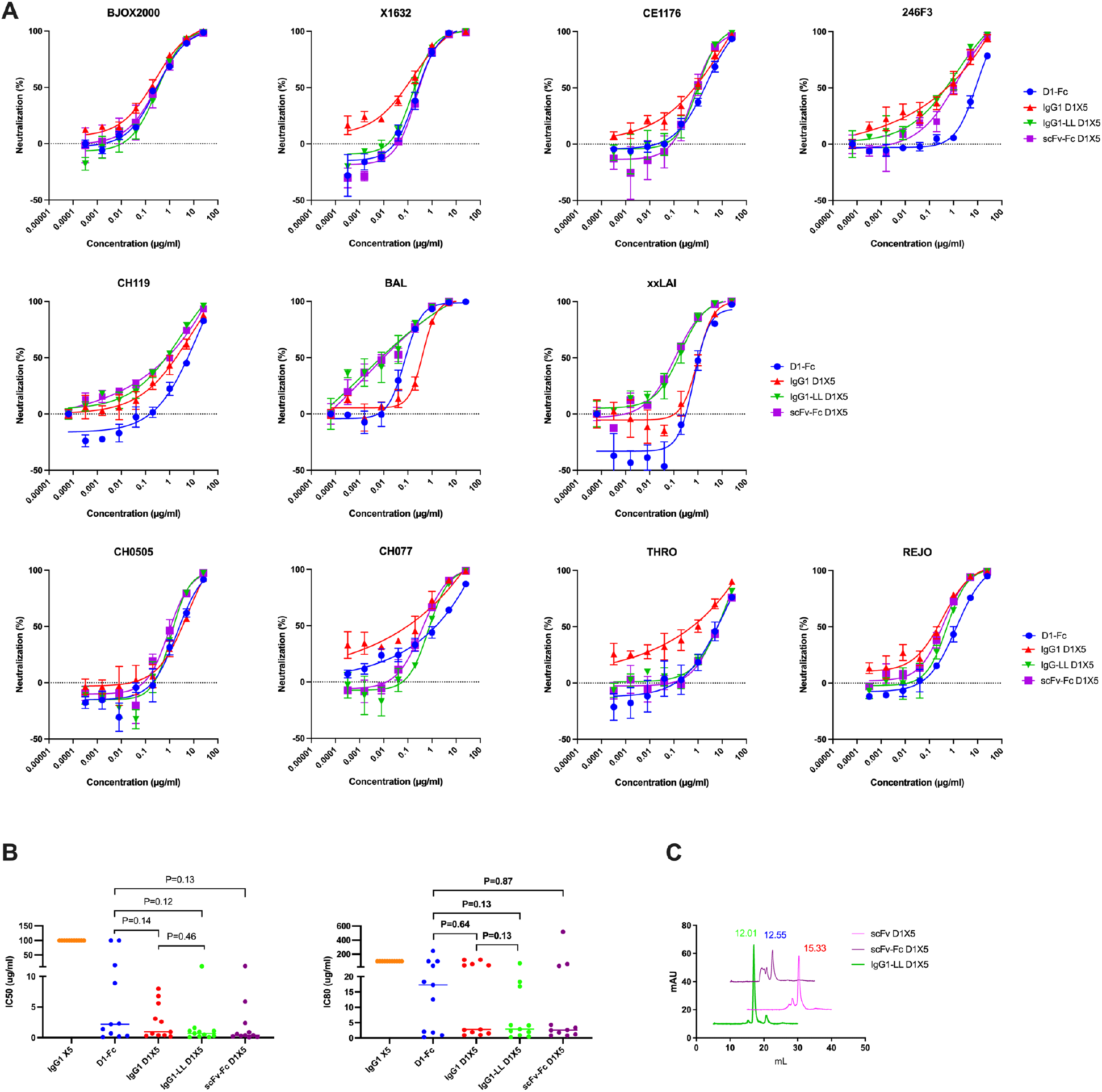
Improved neutralization IC_50_ of the bispecific antibody against HIV-1 variants by extending the linker between D1 and X5. (A)Neutralization effects of D1-Fc, IgG1 D1X5, IgG1-LL D1X5, and scFv-Fc D1X5 against HIV-1 PSV variants and lab-adapted variants. Experiments were performed in triplicate. (B)Median IC_50_ (left) and IC_80_ (right) of IgG1 X5, D1-Fc, IgG1 D1X5, IgG1-LL D1X5, and scFv-Fc D1X5 against HIV-1 variants. Significance was tested using unpaired t test. (C) Aggregation evaluation of scFv D1X5, scFv-Fc D1X5, and IgG1-LL D1X5 measured by SEC.

Moreover, compared with D1-Fc, IgG1-LL D1X5 showed a substantial ~4-folds improvement in neutralization potency (IC_50_: 0.6 vs. 2.6 μg/mL; IC_80_: 3.4 vs. 11.2 μg/mL). When converted to molar units, the enhancement reached ~10-folds (IC_50_: 3.2 vs. 32.5 nM; IC_80_: 17.9 vs. 140.1 nM) (**Table 2 and Figure 4**). The improvement was largely driven by converting several previously insensitive HIV-1 variants such as CE1176, 246F3, Bal, xxLAI, CH0505 into highly sensitive ones by IgG1-LL D1X5. Although accessed on a limited panel, these results suggested that IgG1-LL D1X5 achieves neutralization potency approaching that of elite bnAbs (IC_50_ of 0.6 μg/mL). A head to head comparison with established bnAb aross a larger panel of HIV-1 variants representing the major globally circulating subtypes will be needed to further profile the neutralization breadth and potency of our CD4-CD4i bispecific IgG1-LL D1X5.

By evaluating aggregation and pharmacokinetics of our lead construct, IgG1-LL D1X5 exhibited a predominantly monomeric species by size-exclusion chromatography, whereas scFv-Fc D1X5 displayed both monomeric and polymeric folding (**Figure 4C**). In addition, IgG1-LL D1X5 exhibited a half-life comparable to that of IgG1 D1X5 and D1-Fc (**Figure S4**).

## Discussion

CD4-CD4i antibodies are designed to enhance HIV-1 neutralization by simultaneously blocking the CD4 binding region and the CD4-induced coreceptor binding site on the HIV-1 gp120 envelope spike[23]. These antibodies are expected to exhibit superior neutralization potency against HIV compared to their individual components by a cooperative binding mechanism. The dual engagement not only increase the association rate constant for HIV binding, but also slows the dissociation rate constant through CD4i component, either by stabilizing the CD4-gp120 complex or by facilitating CD4 rebinding following transient dissociation [23].

Although the concept of combining sCD4 with a CD4i antibody is appealing, generating sCD4-CD4i bispecific antibodies capable of simultaneously engaging CD4bs and CD4-induced epitope (CoRbs) is not straightforward. First, multiple sCD4 molecules may be required to fully expose the CD4i epitope[14]. Second, the kinetics of Env conformational changes following sCD4 binding can intrinsically limit synergy: CD4i antibodies bind weakly to Env in the absence of CD4, and once sCD4 dissociates, Env rapidly reverts to a conformation that re-masks the CD4i epitope, leading to dissociation of the entire sCD4-CD4i complex. Third, the spatial geometry and steric constraints surrounding the CD4bs and CoRbs including V1/V2 domain, V3 loop and associated glycans further complicate the design of an effective CD4-CD4i bispecific.

The Bjorkman group observed only modest synergy between sCD4-D1D2 and a CD4i antibody when sCD4 was fused to the N terminus of the heavy chain of the CD4i antibody E51 using a nine repeat (G_4_S) linker[23]. Our group previously generated an sCD4-CD4i construct consisting of m36.4 (VH sdAb) fused to mD1.22 through a naturally stable and flexible polypeptide linker derived from the M13 bacteriophage pIII capsid protein. However, this design yielded only 2 to 3-fold improvement in neutralization compared with CD4-Ig[53]. A more successful example came from the Farzan group, which used an 11 amino acids or longer linker and a CD4i to CD4 orientation (CoR mimetics to CD4 mimetics) to generate a potent bifunctional molecule capable of simultaneously and cooperatively engaging the CD4bs and CoRbs on a single gp120 monomer[54]. Collectively, these findings highlight that precise control over molecular orientation, domain size, and linker architecture is crucial for achieving effective dual targeting and maximal potency in sCD4-CD4i reagents.

In this study, we designed a prototypic bispecific CD4-CD4i antibody, IgG1 D1X5, and evaluated its neutralization potency using TZM-bl cells. The bispecific IgG1 D1X5 antibody exhibited broad neutralization activity against 11 HIV-1 variants. However, its overall potency was moderate. Although enhanced neutralization was observed for several strains, its efficacy was reduced against others including CE1176, Bal, xxLAI, and CH0505 relative to D1-Fc alone (**Table 2**). To improve the neutralization potency of D1X5, we leveraged structure-based molecular optimization to enhance dual binding of HIV Env.

The previous study demonstrated that the smaller size of X5 in the scFv format compared with IgG format, enhances neutralization efficacy primarily by alleviating steric hindrance[32]. After cell surface CD4 binds to the virus during the post-attachment phase, the space between the virus and the target cell surface is 85 Å, which is insufficient to accommodate the 115 Å IgG antibody but sufficient for antibody fragments like scFv, which are only 40 Å in size [32]. This spatial restriction imposed by the host-cell surface and the virion can be bypassed by the CD4-CD4i bispecific described here, in which CD4 engagement of Env exposes the CD4i epitopes and *in cis* facilitates X5 binding. However, the X5 epitope on the bridging sheet (β2-β3-β20-β21) is surrounded by the V3 base and the V1V2 loops, creating steric hindrance that may limit simultaneous binding of sCD4 and X5. Indeed, our structural modeling suggested that the short (G_4_S)_3_ linker between D1 and X5 may impose spatial constraints that limit the flexibility of the D1 and X5 arms in the bispecific antibody during the dynamic dual binding to gp120, potentially restricting their binding stability. In contrast, the longer (G_4_S)_7_ linker permits greater spatial displacement and flexibility between the D1 and X5 domains during binding (**Figure 3B**).

Additionally, the long-linker format exhibited lower binding energy than the (G_4_S)_3_ short linker counterpart as evaluated by the MM/GBSA method.

The neutralization results aligned with our hypothesis, demonstrating that both IgG1-LL D1X5 and scFv-Fc D1X5 exhibited enhanced neutralization against the HIV variants compared to IgG1 D1X5. The lead IgG1-LL D1X5 shows ~ 10 folds neutralization enhancement compared to D1-Fc alone.

The neutralization potency of IgG1-LL D1X5 varied across the panel, with IC_50_ values ranging from 0.08 to 11.5 μg/mL and IC_80_ values from 0.3 to 71.7 μg/mL. This differential sensitivity to sCD4-CD4i bispecific may reflect be variable steric hindrance imposed by the V1/V2/V3 loops and surrounding glycans at the CD4bs and CoRbs, which can restrict access of both sCD4 and CD4i antibodies[55]. Alternatively, intrinsic differences in Env triggerability following CD4 engagement may influence the extent of conformational opening and CD4i epitope exposure, thereby affecting the synergy between sCD4 and CD4i binding[56]. Structural and mutagenesis studies will be important to elucidate the mechanism underlying this variable neutralization sensitivity. In addition, evaluation against a broader panel of HIV-1 variants, alongside comparison with established bnAbs, is needed to more fully define the neutralization breadth of IgG1-LL D1X5.

## Author contribution

WL, XC, JWM, DSD conceived and designed the research; XC designed and characterized all the formats of bispecific antibodies, and performed pharmacokinetic assays; CS, KJP performed neutralization assay; ZS, ZF analyzed binding energy by in silico methods; CP characterized the purity of antibodies. XC wrote the draft of the article, and ZS, WL, KJP, DSD, and JWM revised the manuscript. All authors approved the final manuscript.

## Funding

This work was supported by NIAID funding “REACH: Research Enterprise to Advance a Cure for HIV” (UM1AI164565) and I4C 2.0: Immunotherapy for Cure (1UM1AI164556).

## Acknowledgments

We would like to thank the members of the Center for Antibody Therapeutics for their helpful discussions, and support from the Rustbelt Center for AIDS Research (P30 AI036219) and Research Enterprise to Advance a Cure for HIV (REACH). We thank the University of Pittsburgh Center for Research Computing for molecular dynamic simulation of CD4-CD4i binding to HIV gp120.

## Conflict of interest statement

D.S.D. designed this study while affiliated with University of Pittsburgh School of Medicine. J.W.M. is a consultant to Gilead Sciences, and receives compensation from Galapagos, NV (unrelated to the current work).

## Data availability

The original contributions presented in the study are included in the article/supplementary material. Further inquiries can be directed to the corresponding authors.

## Notes

### Competing Interest Statement

The authors have declared no competing interest.

