## Supplementary data for "Potent and Broad HIV-1 Neutralization by a Bispecific CD4-CD4i Fusion Protein based on Single-Domain CD4-D1 and X5 CD4i antibody"

### Bi-specific anti-HIV antibody (biAb) showed broad neutralization efficacy against HIV variants

**Supplementary Table 1. Neutralization ($IC_{50}$) of bispecific antibody with scFv format against HIV variants**

| HIV Variants | Tier | Country of origin | scFv X5 | scFv D1X5 | scFv-Fc D1X5 |
|---|---|---|---|---|---|
| | | | HIV PSV[1] | | |
| BJOX2000 | 2 | China | 20.8 | 0.7 | 0.3 |
| X1632 | 2 | Spain | 13.4 | 0.1 | 0.2 |
| CE1176 | 2 | Malawi | 6.3 | 4.5 | 0.6 |
| 246F3 | 2 | Tanzania | 3.1 | 1.4 | 2.4 |

| | | | | | |
|---|---|---|---|---|---|
| CH119 | 2 | China | >100 | >100 | 12.2 |
| | | | Lab-adapted strains | | |
| Bal | 1b | US | >100 | 0.1 | 0.04 |
| xxLAI | 1a | France | >100 | 0.1 | 0.1 |

[1]PSV: Pseudovirus; $IC_{50}$ were in µg/mL.

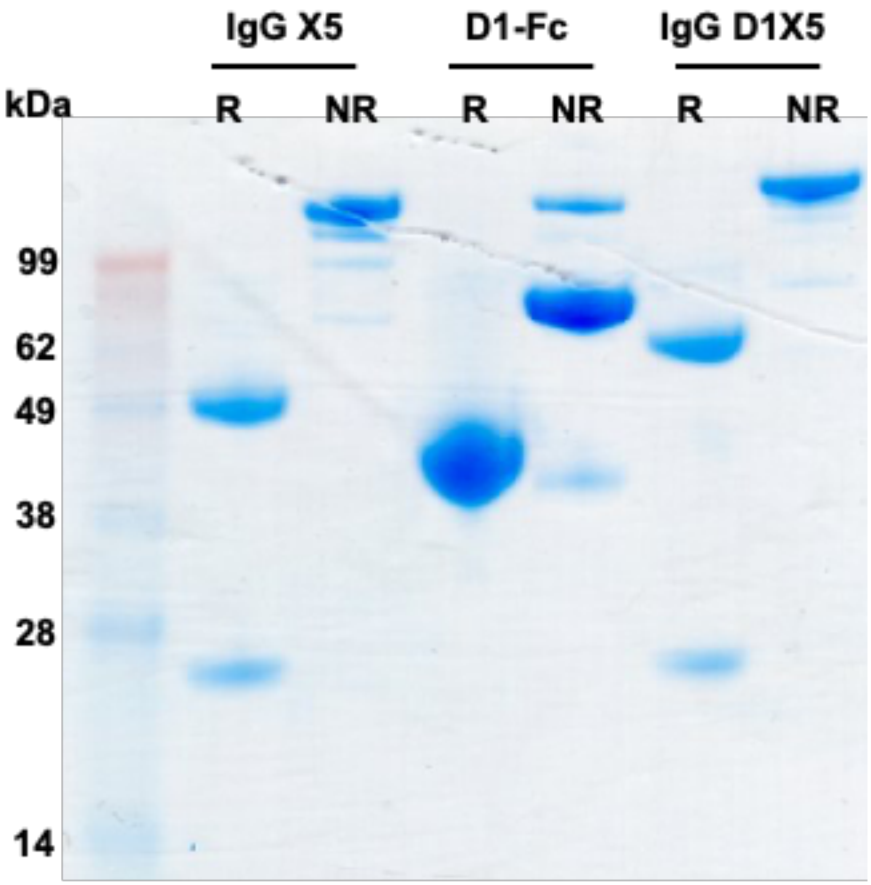


**Supplementary Figure 1: The purity of IgG1 X5, D1-Fc, and IgG1 D1X5 tested by SDS-PAGE.** NR: Non-Reducing; R: Reducing.

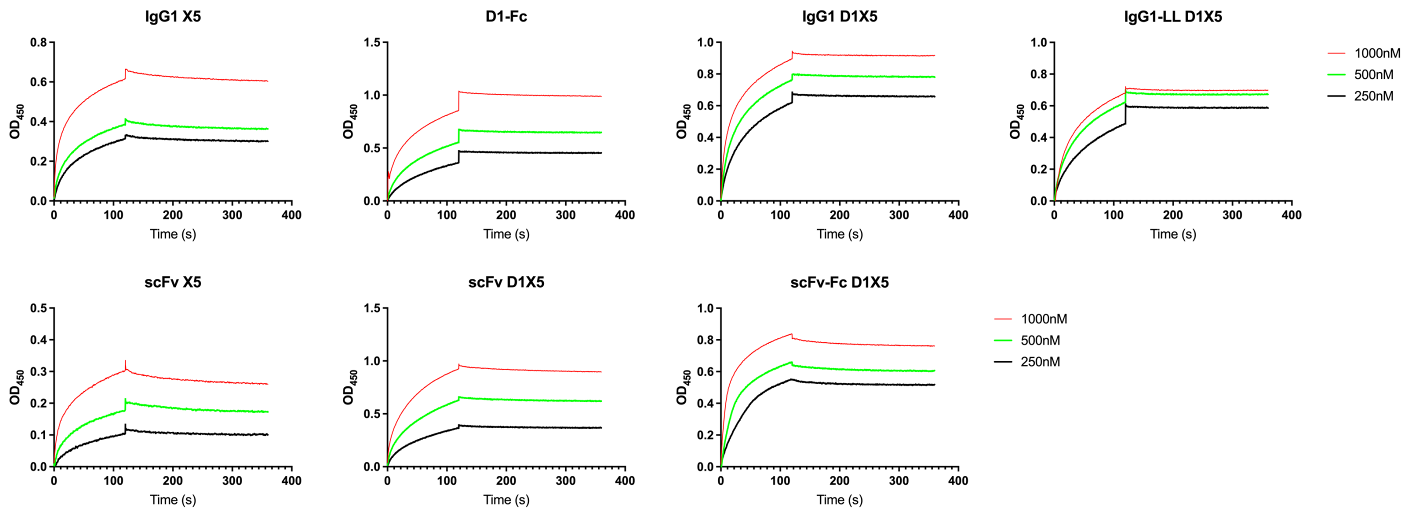


**Supplementary Figure 2: Affinity and avidity characterization of different format of bispecific Ab D1X5 by Blitz.** Upper layer: Kinetics of IgG1 X5 alone, D1-Fc alone, IgG1 D1X5 with short liker $(G_4S)_3$ and IgG1 D1X5 with long liker $(G_4S)_7$; Lower layer: Kinetics of scFv X5 alone, scFv D1X5, and scFv-Fc D1X5 with long liker $(G_4S)_7$.

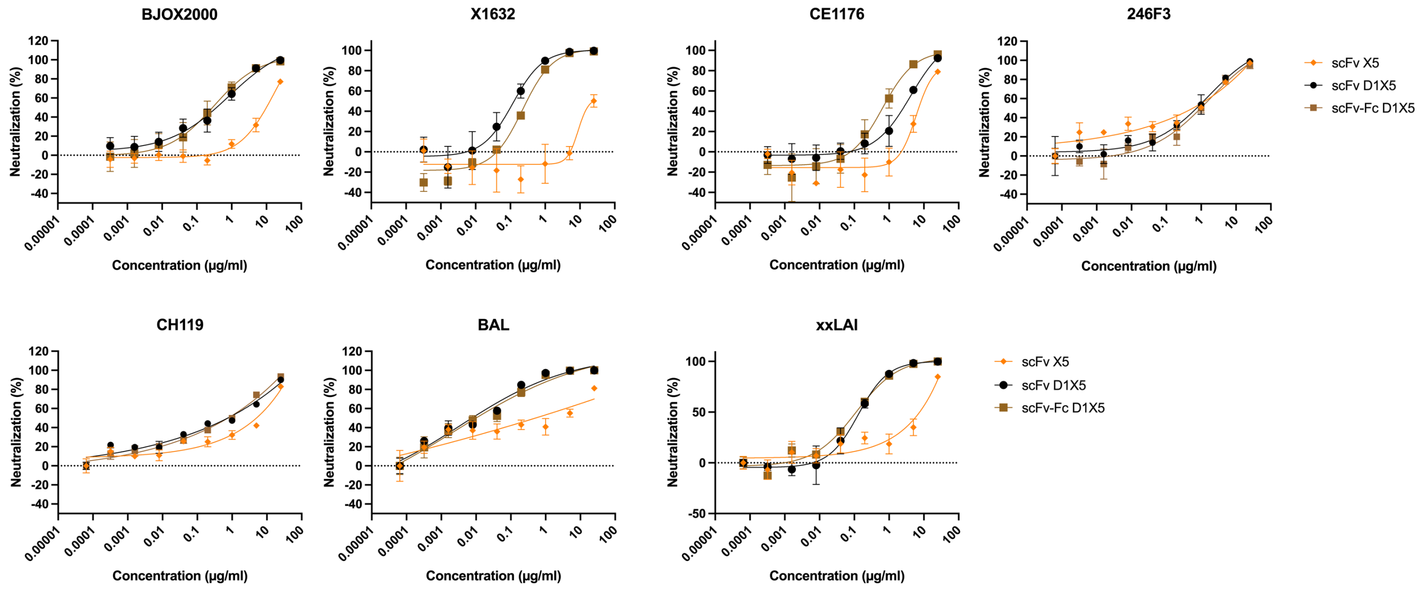


**Supplementary Figure 3: Neutralization of scFv X5 alone, scFv D1X5, and scFv-Fc D1X5 with long liker $(G_4S)_7$ against HIV-1 PSV variants and lab-adaptive variants.** PSV variants: BJOX2000, X1632, CE1176, 246F3, and CH119; Lab-adaptive variants: BAL and xxLAI.

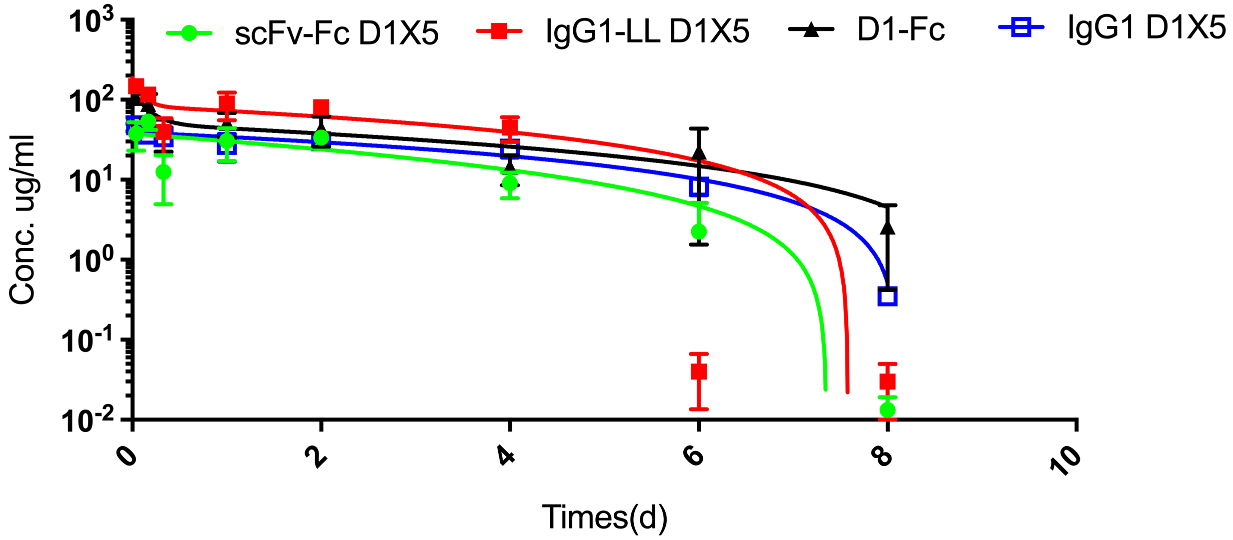


**Supplementary Figure 4: In vivo pharmacokinetics (PK) test.**

PK of scFv-Fc D1X5, IgG1-LL D1X5, IgG1 D1X5, and D1-Fc were tested in BALB/c mice (n=3). Blood was collected at indicated time points and antibody concentration in serum were tested by indirect ELISA.
